# Molecular Dynamics Simulations Unveil the Aggregation Patterns and Salting out of Polyarginines at Zwitterionic POPC Bilayers in Solutions of Various Ionic Strengths

**DOI:** 10.1101/2024.04.24.590968

**Authors:** Man Nguyen Thi Hong, Mario Vazdar

## Abstract

This study employs molecular dynamics (MD) simulations to investigate the adsorption and aggregation behavior of simple polyarginine cell-penetrating peptides (CPPs), specifically modeled as R_9_ peptides, at zwitterionic phosphocholine POPC membranes under varying ionic strengths of two peptide concentrations and two concentrations of NaCl and CaCl_2_. The results reveal an intriguing phenomenon of R_9_ aggregation at the membrane, which is dependent on the ionic strength indicating a salting-out effect. As the peptide concentration and ionic strength increase, peptide aggregation also increases, with aggregate lifetimes and sizes showing a corresponding rise, accompanied by the total decrease of adsorbed peptides at the membrane surface. Notably, in high ionic strength environments, large R_9_ aggregates, such as octamers, are also observed occasionally. The salting-out, typically uncommon for short positively charged peptides, is attributed to the unique properties of arginine amino acid, specifically by its side chain containing amphiphilic guanidinium (Gdm^+^) ion which makes both intermolecular hydrophobic like-charge Gdm^+^ – Gdm^+^ and salt-bridge Gdm+ – C-terminus interactions, where the former are increased with the ionic strength, and the latter decreased due to electrostatic screening. The aggregation behavior of R_9_ peptides at membranes can also linked to their CPP translocation properties, suggesting that aggregation may aid in translocation across cellular membranes.

## Introduction

Peptide adsorption at the cellular membrane is a first and critical prerequisite for their successful subsequent translocation to the cell.^1^ Cellular membranes are highly complex and are composed of many different lipids forming two separate leaflets together with embedded or associated proteins, thus making a very heterogeneous media but with a strongly hydrophobic interior originating from hydrocarbon lipid tails.^2,3^ This makes it difficult for charged species to be efficiently translocated, due to the huge energy penalty of carrying charged ions across a hydrophobic interior.^4,5^ Therefore, in normal, active cellular-controlled processes, endocytosis (or exocytosis) which is driven by ATP and catalyzed by the membrane protein machinery, represents the main pathway to the cellular interior of any cargo to the cells but is often not very cargo-specific.^6,7^

Remarkably, it has been shown in the past that some charged peptides, often rich in positively charged residues, translocate the cellular membranes as efficiently, and do not need ATP as an energy source. In particular, so-called cell-penetrating peptides (CPPs) with a typical length of 5 – 30 amino acids are often composed of a large ratio of arginine (Arg) and lysine (Lys).^8–10^ Interestingly, Arg is more often found in the CPP sequences than Lys and homo-oligomers of Arg peptides in the length of 5 – 15 units show superior translocation compared to their equally charged Lys counterparts, often referred to as arginine “magic”.^11^ The reason behind their larger propensity is not completely clear from the molecular point of view and the details of the translocation mechanism are not understood quantitatively. This stems from the fact that the typical procedures for the determination of free energy of translocation along the *z*-axis of the membrane bilayer are not optimal and result in too high translocation barriers,^12^ suggesting that another more efficient translocation pathway is operative instead. Recently, new reaction coordinate protocols coupled with transient pore formation, have been suggested to obtain the energetics of the process.^13–16^ However, the energetic estimation of charged peptide translocation is still far from being reliably quantified.

In our previous work, we have established by molecular dynamics (MD) simulations that oligoarginines (such as nona-arginines, R_9_) efficiently adsorb to zwitterionic phosphocholine (PC) membranes with free energy of adsorption being about 5 kcal mol^-1^, in contrast to nona-lysines (K_9_) which are not adsorbed to PC membranes at all.^17–21^ Even though both R_9_ and K_9_ have the same positive charge, we have shown by MD simulations that R_9_ can interact with the PC phosphate group in the membrane interior more efficiently than K_9_. Additionally, in the case of R_9_ adsorption, a stronger hydrophobic effect also contributes to the final energy balance.^21^ Moreover, several experimental works from our groups have shown that the adsorption of oligoarginines at PC bilayers is not a computational artifact and is indeed occurring using various experimental techniques, such as lipid/polydiacetylene colorimetric assays^22^ or fluorescence assays on supported lipid bilayers.^19^ It is important to note that zwitterionic, but overall neutral PC bilayers have been chosen as a model lipid membrane since they constitute a majority of lipids in the outer cellular leaflet, where peptide adsorption occurs upon translocation to the cell from the outside.^23^ We have also shown by both MD simulations and fluorescence experiments that ionic strength is an important player in peptide adsorption strength.^20^ Specifically, we have computationally predicted and experimentally shown that by increasing either salt or peptide concentration, the adsorption strength weakens due to an increase in ionic strength which screens electrostatic interactions between the involved charged peptide and lipid species.

However, another interesting phenomenon observed in MD simulations is the attraction and aggregation of like-charged R_10_ and R_9_ (vs K_10_ and K_9_) peptides in water and at membranes, respectively, which is counterintuitive from the electrostatic point of view. In addition to MD simulations which robustly show the Arg aggregation in both media, the aggregation of R_10_ (and lack of K_10_ aggregation) in aqueous solutions has been experimentally suggested by SAXS measurements.^24^ Similarly, the aggregation of R_9_ (but not K_9_) at membranes has been indicated by fluorescence assay experiments at supported lipid PC bilayers.^19^ The reasons behind the “magical” pairing of Arg-rich peptides can be attributed to the chemical nature of the guanidinium cation (Gdm^+^), found in the side chain of Arg amino acid. Our previous findings by extensive MD simulations at both classical^25,26^ and the *ab initio* level,^25,27,28^ indicate an unexpected counterintuitive self-aggregation tendency of Gdm^+^ cations in water, despite their positive charge due to attractive van der Waals interaction originating from specific Gdm^+^ geometry and charge distribution which overcome repulsive electrostatic interaction between positively charged cations. This behavior is not observed in equally charged spherical ammonium (NH_4_^+^) ions, which hints at why Lys-rich peptides do not show aggregation ability.

The tight adsorption of Arg-rich peptides to PC bilayers has been quantified in our previous simulations and experiments,^19,21^ but the aggregation propensity of Arg-rich peptides at the membrane surface remains enigmatic. To resolve this puzzle, in this work, we focus on the detailed molecular view of the R_9_ aggregation propensity at 1-palmitoyl-2-oleoyl-sn-glycero-3-phosphocholine (POPC) using available unbiased microsecond MD simulations and try to assess it in a more detailed and quantifiable way. As our workhorse system, we analyzed simulations of aggregation of R_9_ at POPC bilayers in solutions of different ionic strengths of NaCl and CaCl_2_ using the scaled-charge approach using the ProsECCo force field which corrects for the overstabilization of electrostatic interactions in non-polarizable simulations.^29–31^ This force field has been successfully applied in many systems, ranging from ions in aqueous solutions to the adsorption of ions at phospholipid membranes.^32–36^ Specifically, we concentrate on the quantification of aggregation propensity and aggregation patterns of R_9_ peptides at membranes in different peptide and salt concentration conditions in molecular detail, which is relevant for further studies of the CPP penetration mechanism which has still not been connected to their increased translocation ability.^17^

### Computational Methods

Atomistic molecular dynamics (MD) simulations were conducted to examine the interactions of nona-arginines (R_9_) in aqueous solutions of either NaCl or CaCl_2_ with POPC bilayers in our previous work.^20^ Whereas the focus of the former analysis was to assess the adsorption energetics, here we focused on the aggregation properties of R_9_ at membranes. We analyzed nine simulations performed for two concentrations of R_9_ peptides (0.021 and 0.056 m) and two concentrations of NaCl (0.133 m, and 1.065 m) or CaCl_2_ salts (0.133 m, and 1.065 m), as well as the reference simulation without added salt, respectively. Initially, peptides (6 or 16 R_9_ molecules corresponding to the concentrations above) were evenly distributed in the bulk phase on both sides of the POPC bilayer. Chloride counterions are used to neutralize the systems, and the simulation boxes for all studied systems contained 15,856 water molecules. Additional NaCl and CaCl_2_ salts were added in the concentrations described above. The membrane bilayer comprised 100 1-palmitoyl-2-oleoyl-sn-glycero-3-phosphocholine (POPC) lipids in each leaflet, resulting in a total of 200 lipids.

This study employed CHARMM36m-based ProsECCo models for lipids and peptides,^29^ along with an electronic continuum correction (ECC) approach for ion parameters to address the overbinding issue of charged molecules to zwitterionic bilayers.^30,31,37^ Specifically, partial charges in the ProsECCo models, including those of the phosphate and choline groups of POPC, peptide termini, and the charged groups of Arg, were adjusted to scale down the total charge of each group from +1 to +0.75 (Table S1). No other modifications were made to the CHARMM36m lipid or peptide parameters.^38^ NBFIX was disabled, as the ECC approach achieved a similar effect without the need for an additional *ad hoc* correction.^29^ All used topologies are available in GitLab at https://gitlab.com/sparkly/prosecco/prosECCo75. The systems were solvated with CHARMM-specific TIP3P (“TIPS3P”) water,^39^ and buffered Verlet lists were employed for tracking atomic neighbors with a 1.2 nm cut-off for the Lennard-Jones potential. Van der Walls interactions were treated using a cut-off of 1.2 nm, with the forces smoothly attenuated to zero between 1.0 and 1.2 nm. Long-range electrostatics were treated using the smooth particle mesh Ewald method.^40^ After the steepest descent minimization of all systems, for the equilibration we used the Berendsen thermostat and barostat,^41^ while the production runs utilized the Parrinello–Rahman barostat^42^ with a semi-isotropic pressure coupling and a 1 bar reference pressure. The Nosé–Hoover thermostat^43^ was applied with a target temperature of 310 K. Covalent bonds involving hydrogens in peptides and lipids were constrained using the P-LINCS algorithm,^44^ and the SETTLE algorithm was used for constraints in water molecules.^45^ MD simulations were conducted for 2 μs with a time step of 2 fs using the GROMACS package.^46^ Analysis of production trajectories was performed using in-house Python scripts in conjunction with the MDAnalysis library,^47^ omitting the first 500 ns from the analyses, while VMD was used for the generation of MD simulations figures.^48^

## Results and Discussion

## Number density profiles and R_9_ aggregation propensity at POPC bilayer

First, we summarize the analysis of the peptide adsorption reported previously.^20^ The symmetrized number density profiles of R_9_ adsorption to the POPC membrane, for two different peptide concentrations, i. e. low (0.021 m) and high (0.056 m) in different salt concentrations (Figure 1, top and bottom panels). The non-symmetrized number density profiles are shown in Figure S1 indicating a good qualitative agreement with symmetrized ones (apart from inevitable quantitative differences due to thermal equilibrium) and proving that the simulation time is sufficient for proper sampling. The number of adsorbed peptides decreases with both peptide and salt concentration, which is more pronounced in the case of CaCl_2_ due to the higher ionic strength of the solution. Interestingly, in the case of the high peptide concentration and the highest CaCl_2_ concentration (Table 2), the adsorption of R_9_ is quite weak (but still existing), which can be explained by the decrease of the electrostatic interactions between positively charged peptides and negatively charged POPC phosphate groups due to high ionic strength.^20^

**Fig. 1.**
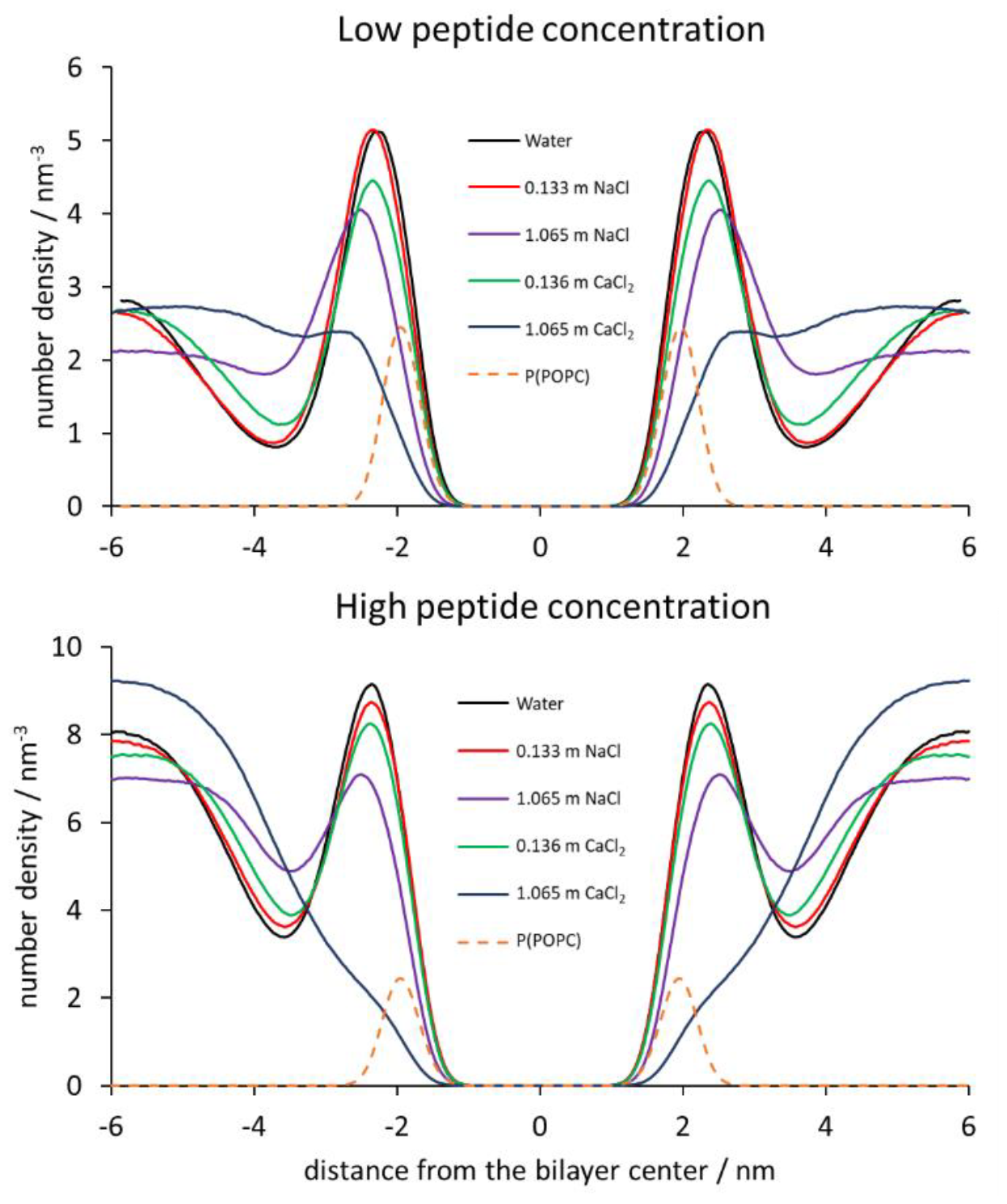
Symmetrized number density profiles for peptide center of mass for low R_9_ concentration (upper panel) and high R_9_ concentrations (bottom panel) in different systems with respect to the distance from the POPC bilayer center. The number density of POPC phosphorus atoms is shown as the dashed line for the reference system without added salt for low and high peptide concentrations.

Using number density profiles only, it is not possible to assess the degree of peptide aggregation since the profiles are time-averaged in the analysis and mask possible aggregation events. Here, we assessed the aggregation propensity by analyzing the individual MD simulation frames and checking whether in the vicinity of adsorbed peptides at POPC other peptides are found (not necessarily bound to POPC), using a cut-off of 0.6 nm between any atoms of the peptide in contact. If the criterion is fulfilled, we denote the associated peptides as an aggregate. Tables 1 and 2 show the calculated total degree of aggregation (*i. e*. the number of frames with some aggregate form of any size divided by the total analyzed frames), for low and high peptide concentrations. In addition, for each of the aggregates (regardless of its degree of oligomerization which is discussed later), we analyzed the contribution of different interactions between selected groups in R_9_ (Gdm^+^ – Gdm^+^, Gdm^+^ – COO^-^ and NH_3_^+^ – COO^-^) given as a ratio of a number of contacts between two given groups (also with a cut-off of 0.6 nm) vs. the total number of three selected contact types within this threshold distance.

**Table 1.**
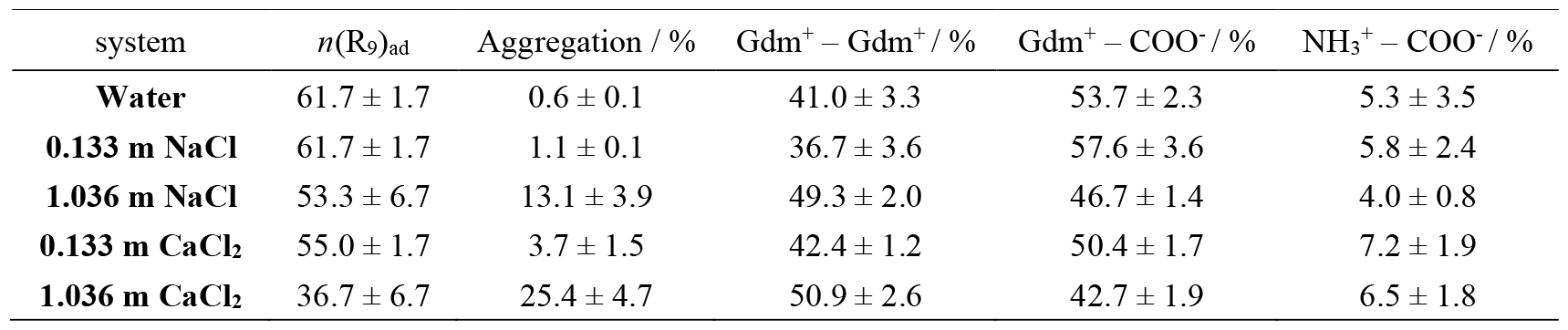
The probability of adsorption of R_9_ peptides at POPC, *n*(R_9_)_ad_, and R_9_ aggregation events, together with the ratio of different group interactions is given in percentages in systems with low peptide concentration. Error bars were estimated by the standard deviation of three individual 500 ns MD simulation blocks.

| system | $n(R_9)_{ad}$ | Aggregation / % | Gdm<sup>+</sup> – Gdm<sup>+</sup> / % | Gdm<sup>+</sup> – COO<sup>-</sup> / % | NH<sub>3</sub><sup>+</sup> – COO<sup>-</sup> / % |
| --- | --- | --- | --- | --- | --- |
| **Water** | 61.7 ± 1.7 | 0.6 ± 0.1 | 41.0 ± 3.3 | 53.7 ± 2.3 | 5.3 ± 3.5 |
| **0.133 m NaCl** | 61.7 ± 1.7 | 1.1 ± 0.1 | 36.7 ± 3.6 | 57.6 ± 3.6 | 5.8 ± 2.4 |
| **1.036 m NaCl** | 53.3 ± 6.7 | 13.1 ± 3.9 | 49.3 ± 2.0 | 46.7 ± 1.4 | 4.0 ± 0.8 |
| **0.133 m CaCl<sub>2</sub>** | 55.0 ± 1.7 | 3.7 ± 1.5 | 42.4 ± 1.2 | 50.4 ± 1.7 | 7.2 ± 1.9 |
| **1.036 m CaCl<sub>2</sub>** | 36.7 ± 6.7 | 25.4 ± 4.7 | 50.9 ± 2.6 | 42.7 ± 1.9 | 6.5 ± 1.8 |

The analysis of the aggregation data shows several interesting facts. The number of adsorbed R_9_ peptides decreases with the salt concentrations which is more pronounced in the case of CaCl_2_ than NaCl, due to the higher ionic strength of solutions, as shown in our previous work,^20^ and agrees with the number density profiles shown in Figure 1. However, the peptide aggregation propensity is reversed – the higher the ionic strength is, the peptide aggregation increases, ranging from a negligible propensity in neat water solutions at low peptide concentrations (0.6 %, system without added ions, Table 1) up to a significant aggregation (50.4 %, 1.065 m CaCl_2_, Table 2) in high peptide concentration systems with high CaCl_2_ concentration. This resembles a biologically very important phenomenon in protein chemistry, where the solubility of a protein decreases in the presence of high concentrations of salts, known as salting-out.^49,50^ However, it is interesting that salting-out occurs here for highly positively charged Arg oligomers at membranes, which are charged and relatively short. The detailed analysis shows that the amount of like-charge Gdm^+^ – Gdm^+^ interactions, which are mostly hydrophobic,^28^ increase with the salt concentration thus contributing mostly to the increase in peptide aggregation. This is slightly more pronounced for high peptide concentrations where the aggregation is more pronounced too (Tables 1 and 2). On the other hand, the electrostatic interactions between Gdm^+^ – COO^-^ and NH_3_^+^ – COO^-^ decrease with the addition of salt due to charge screening. Importantly, the increase in the peptide aggregation does not lead to a larger number of adsorbed peptides since the peptide aggregation is the strongest in the cases where the least peptides are adsorbed (system with 1.065 m CaCl_2_, Table 2), showing an inverse correlation between peptide adsorption vs. peptide aggregation.

**Table 2.** The probability of adsorption of R_9_ peptides at POPC, *n*(R_9_)_ad_, and R_9_ aggregation events, together with the ratio of different group interactions is given in percentages in systems with high peptide concentration. Error bars were estimated by the standard deviation of three individual 500 ns MD simulation blocks.

| system | $n(R_9)_{ad}$ | Aggregation / % | Gdm<sup>+</sup> – Gdm<sup>+</sup> / % | Gdm<sup>+</sup> – COO<sup>-</sup> / % | NH<sub>3</sub><sup>+</sup> – COO<sup>-</sup> / % |
| --- | --- | --- | --- | --- | --- |
| **Water** | 45.6 ± 0.6 | 13.9 ± 1.7 | 42.3 ± 0.9 | 51.3 ± 0.7 | 6.4 ± 0.8 |
| **0.133 m NaCl** | 44.4 ± 0.6 | 18.0 ± 3.1 | 44.6 ± 0.5 | 49.7 ± 0.6 | 5.6 ± 0.7 |
| **1.065 m NaCl** | 38.1 ± 2.5 | 32.2 ± 3.5 | 50.7 ± 0.6 | 45.5 ± 0.6 | 3.8 ± 0.4 |
| **0.133 m CaCl<sub>2</sub>** | 41.9 ± 1.3 | 16.1 ± 0.6 | 45.3 ± 0.5 | 48.5 ± 0.5 | 6.2 ± 0.5 |
| **1.065 m CaCl<sub>2</sub>** | 16.9 ± 1.9 | 50.4 ± 5.0 | 53.9 ± 1.3 | 42.5 ± 1.6 | 3.6 ± 0.7 |

### The diversity and lifetime of adsorbed R_9_ aggregates at POPC

As the next step, we analyzed the distribution of aggregates depending on their size. Figure 2 shows the aggregate distribution for two R_9_ peptide concentrations in different salt solutions. The upper panel shows the distribution for lower peptide concentrations. We see that the amount of dimers is dominating in all systems, being in the range of 80 – 100 %. Only in systems with higher ionic strength (especially in CaCl_2_ solutions), a significant percentage of trimers (< 20%), a small number of tetramers (< 5%), and a very small number of pentamers (< 0.5%, not shown in the Figure) are also present.

**Fig. 2.**
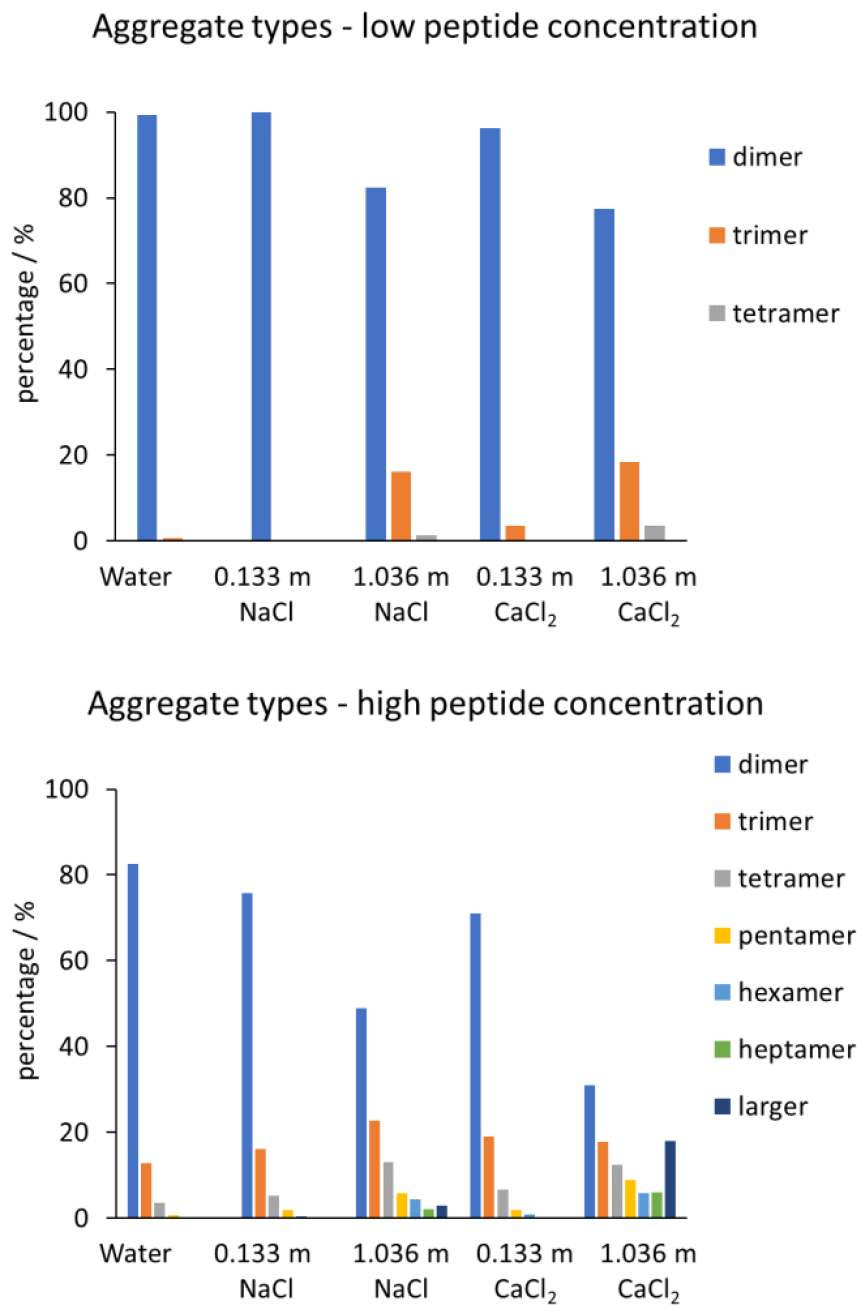
The percentage of different-sized aggregates in systems with low peptide concentration (upper panel) and high peptide concentrations (bottom panel) in systems with different ionic strengths.

A more interesting picture occurs in systems with higher peptide concentrations (bottom panel). Although the dimers are still dominantly present in all systems, the distribution of trimers and higher aggregates is qualitatively different than in the low peptide concentration regime. In particular, in the systems with high ionic strength with 1.065 m NaCl or CaCl_2_, large aggregates also occur in non-negligible amounts. Even more interestingly, in the system with 1.065 m CaCl_2_, the percentage of larger aggregates than heptamers is almost 20%. Although this indicates that the aggregation is sensitive to the ionic strength, we should also keep in mind that in this case, the number of peptides that are indeed adsorbed at the POPC interface is the smallest (Table 2).

In general, we can see the following trends in our MD simulations. First, in the systems with low peptide concentration, the maximum size of aggregates is much smaller compared to higher peptide concentrations reaching the maximum size of pentamer in the system with 1.065 m NaCl and CaCl_2_. Second, the diversity of differently-sized aggregates is much more expressed for higher peptide concentrations, where heptamers and even higher aggregates are present, especially at higher salt concentrations and in turn higher ionic strengths.

In Table 3, we analyzed the average lifetime of R_9_ aggregates adsorbed at the membrane, depending on their size. Overall, the lifetime of same-sized aggregates increases upon increasing the peptide and salt concentration, which is most notable for the dimers. In the systems where many different-sized aggregates are present, the lifetime of larger aggregates is, in general, decreasing compared to dimers, reaching minimum lifetimes of ca. 100 ps for very large aggregates. Therefore, despite the lower number of adsorbed R_9_ peptides upon the increase of peptide and salt concentrations, the aggregated peptides that are adsorbed at the POPC interface have an overall larger adsorption lifetime when all aggregate sizes are considered.

**Table 3.** The average lifetime of R_9_ aggregates (in ps) of different sizes (dimer–heptamer and higher aggregates) at the POPC bilayer for low and high peptide concentrations in systems with different salt compositions and concentrations. The error bar is the standard deviation of the mean value when multiple aggregation events occur. For larger aggregates (> heptamer), the range of aggregate lifetimes is presented.

| system | dimer | trimer | tetramer | pentamer | hexamer | heptamer | > heptamer |
| --- | --- | --- | --- | --- | --- | --- | --- |
| **Water** | <sup>1)</sup> 297 ± 38 | – | – | – | – | – | – |
|  | <sup>2)</sup> 799 ± 50 | 335 ± 20 | 270 ± 32 | 207 ± 43 | 150 ± 29 | – | – |
| **0.133 m NaCl** | 492 ± 88 | – | – | – | – | – | – |
|  | 857 ± 60 | 360 ± 22 | 287 ± 25 | 218 ± 23 | 140 ± 21 | 220 ± 58 | – |
| **1.065 m NaCl** | 928 ± 103 | 652 ± 141 | 380 ± 77 | <sup>3)</sup>~300 | – | – | – |
|  | 638 ± 26 | 425 ± 18 | 352 ± 18 | 241 ± 17 | 258 ± 21 | 159 ± 13 | 150 – 200 |
| **0.133 m CaCl<sub>2</sub>** | 760 ± 119 | 231 ± 55 | <sup>3)</sup>~100 | – | – | – | – |
|  | 561 ± 31 | 337 ± 19 | 254 ± 20 | 159 ± 71 | 129 ± 14 | 217 ± 48 | <sup>3)</sup>~100 |
| **1.065 m CaCl<sub>2</sub>** | 1030 ± 144 | 493 ± 53 | 366 ± 65 | 138 ± 18 | – | – | – |
|  | 659 ± 48 | 443 ± 31 | 321 ± 23 | 273 ± 20 | 237 ± 20 | 244 ± 19 | 150 – 250 |

<sup>1)</sup> The upper row shows data for low peptide concentration.
<sup>2)</sup> The lower row shows data for high peptide concentration.
<sup>3)</sup> A single continuous aggregation event.

### Visualization of different aggregates at the POPC interface

Finally, despite the quantitative analysis of the distribution of R_9_ aggregates at POPC, it is instructive to visualize the aggregates at the membrane surface. In Figure 3 we show a selected typical snapshot from the MD simulation of the reference system, where no additional salt is added to the system. According to Table 1, we see that in the system with low peptide concentration, the aggregation of R_9_ is rare, being observed only in less than 1% of the simulation time.

**Fig. 3.**
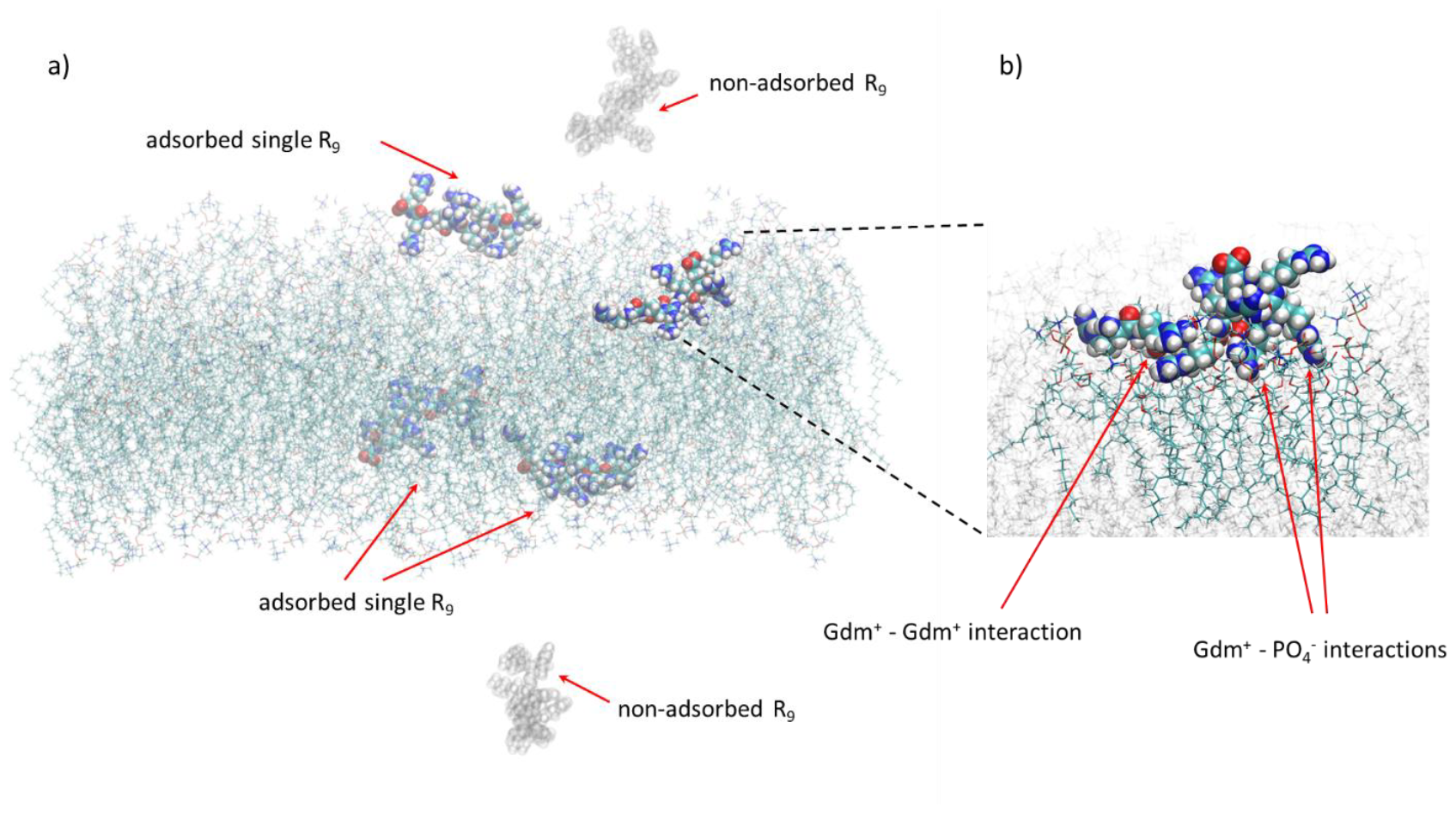
A selected snapshot from MD simulations showing the lack of aggregation of R_9_ peptides in the reference system, without added salt at low peptide concentration. In panel a) we observe four adsorbed R_9_ peptides (opaque representation) and two non-adsorbed R_9_ peptides (ghost representation). Panel b) shows the magnified region where one of the R_9_ peptides is adsorbed at the POPC interface with indicated intramolecular Gdm^+^ – Gdm^+^ interaction and deep penetration of R_9_ sidechains and interaction of its Gdm^+^ groups with POPC phosphate groups.

A detailed analysis of the MD snapshot shows a typical situation when peptides do not aggregate at the POPC surface (Figure 3a). When no aggregation is present, peptides lie down almost parallel to the membrane surface and sidechains are inserted in the membrane interior where Gdm^+^ cations interact strongly with the POPC phosphate groups (Figure 3b). This has been observed in many simulation studies so far, and the detailed analysis of the energetics of this particular interaction is presented in Ref 21. Notably, we also observe the intramolecular like-charge Gdm^+^ – Gdm^+^ interaction (Figure 3b), which is typical for arginine-rich peptides as suggested previously by MD simulations.^19,22,24^

In the case when the concentration of R_9_ peptides is high, together with high CaCl_2_ concentration, the probability of aggregation of R_9_ increases, as suggested in Tables 1 and 2. We see that R_9_ aggregates can be formed in different sizes, as illustrated in Figure 4 (left panel) where we see an R_9_ octamer and R_9_ trimer which are bound to the POPC interface, but do not lie perpendicularly to the membrane surface, like in the case when single R_9_ molecules are adsorbed. Instead, they form a cluster where one peptide (in the case of a trimer) or two peptides (in the case of an octamer) are adsorbed to the interface, and other peptides are aggregated on top of them. For this particular system, this is also reflected in the reduced R_9_ number density (Figure 1) in the vicinity of the membrane together with the calculated number of adsorbed peptides (Table 2), which is significantly lowered compared to systems with lower peptide concentration and ionic strength. Nevertheless, the peptides that are adsorbed (although in a smaller quantity), are in a large ratio adsorbed as aggregates (> 50%, Table 2). The weaker adsorption of the aggregate is not unexpected, since the Gdm^+^ groups, responsible for tight adsorption of single R_9_ to POPC phosphate group, are consumed in the intermolecular interactions between R_9_ peptides in the aggregate and are not available for interaction with POPC membrane (see below).

**Fig. 4.**
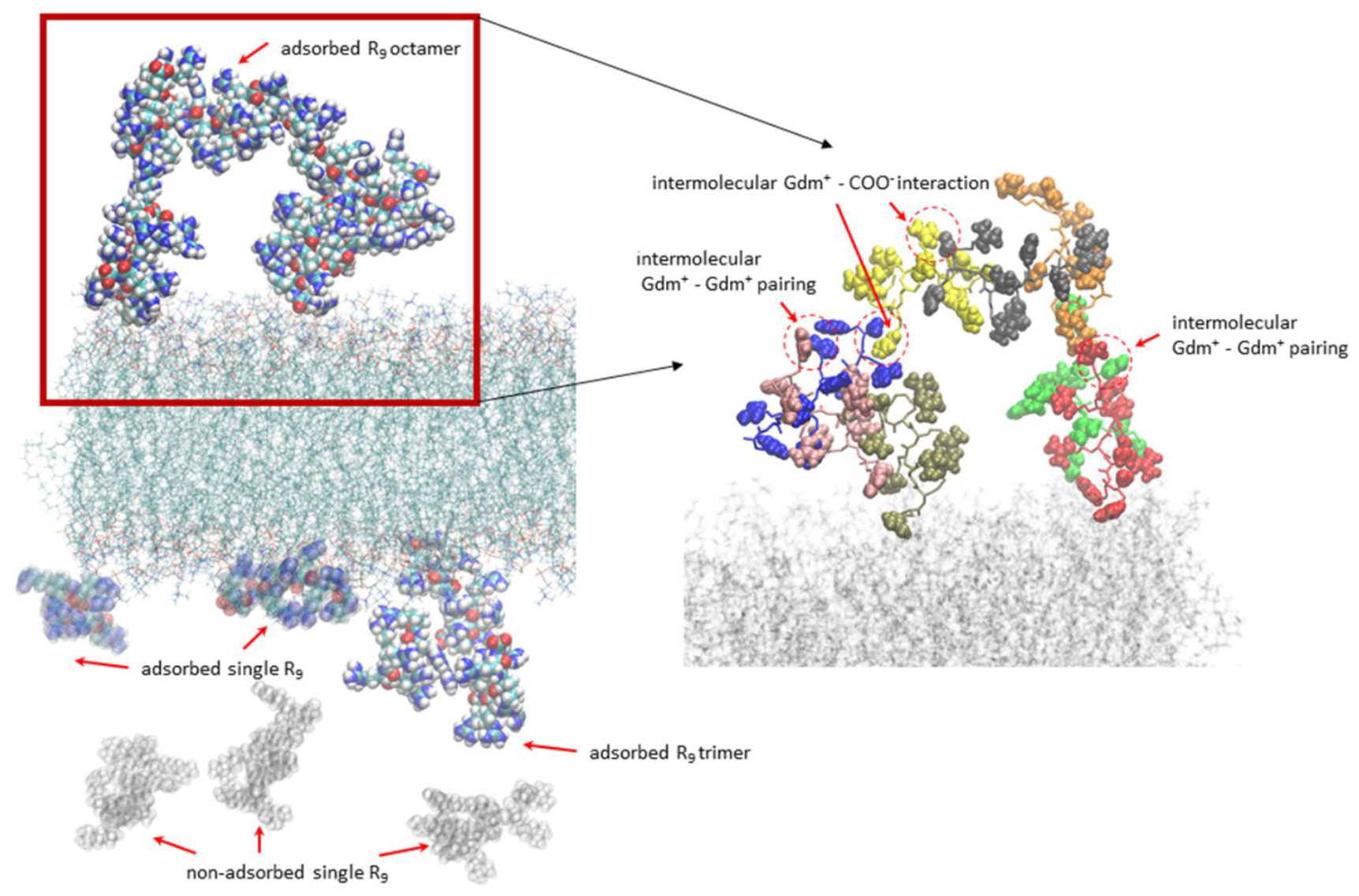
Left panel: A selected snapshot from MD simulations showing the aggregation of R_9_ peptides in the system with 1.065 m CaCl_2_ at high peptide concentration. The R_9_ octamer and R_9_ trimer are shown in the opaque representation. Two single-adsorbed R_9_ peptides are shown in the transparent representation, whereas non-adsorbed R_9_ peptides in the bulk are shown in the ghost representation. Right panel: The selected intermolecular interactions in the adsorbed R_9_ octamer. Individual R_9_ peptides are shown in the licorice representation with different colors, while the corresponding Gdm^+^ and COO^-^ groups are shown in the vdW representation. Gdm^+^ - Gdm^+^ like-charge pairing and salt-bridge-like Gdm^+^ - COO^-^ interactions are indicated with the red arrows and shown in the red dashed circles.

Finally, let’s take a look at one of the aggregates and describe in more detail the interactions, which are quantified in Tables 1 and 2. The intermolecular interactions in the R_9_ octamer are presented in the right panel of Figure 4. We see that the intermolecular aggregate has many contacts between Gdm^+^ groups with either Gdm^+^ groups of another peptide in the aggregate (like-charge pairing) or with carboxylic COO^-^ group (salt-bridges). This is at odds with our previous studies of polyarginine R_10_ aggregate in water where energetically favored salt-bridge interactions between the negatively charged C-terminus and adjacent Gdm^+^ groups dominate compared to like-charge Gdm^+^ – Gdm^+^ interactions, which are hydrophobic and weaker due to the missing attractive electrostatic contribution.^24^ However, upon the interaction of R_9_ with POPC, the ratio of intermolecular Gdm^+^ – Gdm^+^ interactions is either similar or larger vs. Gdm^+^ – COO^-^ interactions (Tables 1 and 2). This effect is due to the peptide interaction with the membrane which limits its conformational freedom compared to the aqueous solution where polyarginine is more flexible and more easily adopts conformations that enable more favored salt-bridge interactions. More interestingly, the ratio of Gdm^+^ – Gdm^+^ interactions increases with the increase of ionic strength which is another manifestation of the salting-out effect at higher salt concentrations where Gdm^+^ – COO^-^ electrostatic interactions are being effectively screened out and hydrophobic Gdm^+^ – Gdm^+^ interactions are unaffected or even increased as computationally predicted for model hydrophobic systems in water.^51^

### Implications to the cell-penetration peptides (CPPs) translocation mechanism

As mentioned in the Introduction, CPPs, such as nona-arginine R_9_, passively translocate across cellular membranes without the need for ATP.^8–10^ This property is especially interesting in light of controlled drug delivery since it enables the delivery of active compounds to the cells without modifying innate translocation cellular mechanisms, such as endocytosis.^52,53^ The details of the translocation mechanism are not currently known at the molecular level, predominantly due to the inability of current computational techniques to properly evaluate the energy barrier for the translocation of CPPs across the hydrophobic lipid bilayer interior which is almost impermeable for charged species, such as polyarginines used in this work. However, the calculated barriers for arginine/polyarginine translocation using the simple *z*-coordinate are similar to the barriers for lysine/polylysine translocation.^54,55^ Therefore, a question arises – why polylysines do not penetrate across the bilayers as well? For neutral membranes, the reason is clear since it has been shown both computationally and experimentally that K_9_ does not adsorb to the neutral POPC bilayers (which serve as a mimic to the neutral outer cellular leaflet^3^), in contrast to R_9_, so it cannot translocate across bilayers if there is no peptide adsorption.^19,21^ This results from the fact that the interaction of Arg side chain – Gdm^+^ ions with the POPC phosphate groups is stronger compared to the analogous Lys side chain –NH_3_^+^ ions interaction due to the planarity of Gdm^+^ ion which more easily interacts with the buried POPC phosphate group.^21^ An alternative idea that has been suggested by MD simulations is that octa-arginine (R_8_) can stabilize the artificially created pore in the neutral DPPC or negatively charged DOPE/DOPS lipid bilayers, in contrast to octa-lysine (K_8_) thus facilitating R_8_ to translocate the bilayer. ^55,56^

However, the arguments above are not fully sufficient for the understanding of the molecular view of the translocation mechanism. Namely, when lipid bilayers are negatively charged, like for example during the endosomal escape,^57,58^ then both polyarginine and polylysine, which are positively charged, can adsorb at the membrane due to attractive electrostatic forces potentially enabling polylysine to cross the bilayer, which has not been confirmed experimentally.^58^ Also, the energy for the spontaneous pore opening is quite high (around 20 kcal mol^-1^ for DPPC),^59^ similar to energy barriers like those for single arginine/lysine translocation,^54^ thus again not explaining how adsorption of octa-arginine can initiate the opening of the pore in the membrane.

Still, in MD simulations the polyarginine aggregates are always present at the membranes, while polylysines are mutually repelled at the membrane surface, regardless of their respective membrane binding affinity. This has been also experimentally confirmed by fluorescence measurements at supporting lipid bilayers.^19^ As previously computationally suggested, the transfer of single Arg,^60^ (or Gdm^+^ cation)^27^ into lipid bilayers is nonadditive and the same energy penalty for crossing the barrier is paid for the transfer of a single Arg or multiple Arg amino acids. Intuitively, we would expect that the same energy barrier would be also present for a single polyarginine peptide, or several polyarginine peptides (that do not repel each other) which might follow the first peptide across the membrane in a concerted manner once the membrane pore is spontaneously created. Therefore, the existence of polyarginine aggregates adsorbed at the lipid membranes can serve in two ways – first, they can destabilize the membrane due to a high positive charge catalyzing the spontaneous transient pore formation in the membrane, similar to the electroporation effect;^61^ second, the aggregates can also serve as a reservoir for additional polyarginine peptides once the membrane translocation process of polyarginine has started. Unfortunately, the computational estimates of what would be the barriers for single vs. aggregated polyarginine translocation, as well as the energetics for peptide aggregate-induced pore formation, are still missing due to inadequate computational techniques.^12–16^ Nevertheless, we believe that the fact that polyarginines indeed aggregate at model membranes as shown in this work, and are sensitive to ionic strength, is interesting enough to be considered as one of the possible ways how to understand the arginine “magic”^17^ and the fact why polyarginine acts as an efficient CPP.

## Conclusions

In this work, we showed by MD simulations that simple polyarginine CPP (modeled as R_9_ peptide) adsorbs to the zwitterionic POPC membrane in an ionic strength-dependent manner. First, we showed that by the increase of R_9_ concentration, as well as the increase in ionic strength (modeled as 0.133 and 1.065 m NaCl and CaCl_2_ solutions, respectively) the total number of adsorbed peptides decreased, in agreement with our previous work.^20^

However, the analysis of the adsorbed peptides at the POPC bilayer revealed an interesting phenomenon of R_9_ aggregation, which is also dependent on the ionic strength. We analyzed in detail how peptide concentration and ionic strength influence peptide aggregation and showed that by the increase of ionic strength, either by the increase of peptide concentration or the presence of Na^+^ and Ca^2+^ salts, aggregation of peptides is also increased. In particular, we show that the diversity of aggregates and the aggregate size also increase with the ionic strength. The majority of aggregates at the POPC bilayer are present in the form of dimers, but large R_9_ aggregates (heptamers and larger) can be sporadically found at the POPC bilayer in systems with high ionic strength which is a clear indication of the salting-out effect, *i. e*. the loss of peptide solubility with the addition of salt, which is unusual for short positively charged peptides. However, since arginine contains amphiphilic Gdm^+^ ion in the side chain with counterintuitive properties, from like-charge pairing and anisotropic solvation, the salting-out effect is a consequence of its hydrophobic properties and screening of electrostatic interactions in high ionic strength solution. We also analyzed the aggregate lifetimes at the membrane, which in general increase with the peptide concentration and ionic strength for the same-sized aggregates.

Finally, the aggregation of R_9_ peptides at membranes can potentially be connected to the CPP translocation properties of polyarginines. Since the R_9_ peptides are aggregated (or at worst non-repelled) at the POPC membrane, during the transfer across spontaneously created transient pores in the bilayers they might translocate in a concerted manner without additional energy penalty, where arginine aggregates serve as a peptide reservoir. This is contrasted to the transfer of the non-CPPs, such as polylysine peptides, which do not aggregate and do not translocate cellular membranes. Still, the molecular details and energetics of the translocation process are not currently available due to high computational demands and improper reaction coordinates for peptide translocation, which is planned to be addressed in future work.

## Supporting Information

Supporting Information associated with this paper can be found in the online version at http://

## Acknowledgments

M.N.T.H. acknowledges the Faculty of Mathematics and Physics of the Charles University (Prague, Czech Republic) where she is enrolled as a Ph.D. student, and the International Max Planck Research School for “Many-Particle Systems in Structured Environments” (Dresden, Germany) for support. M. V. acknowledges support by the Ministry of Education, Youth and Sports of the Czech Republic through the e-INFRA CZ (ID:90254), Project OPEN-28-18. The authors would like to acknowledge the contribution of COST Action CA21169, supported by COST (European Cooperation in Science and Technology). We also thank Prof. Pavel Jungwirth and Dr. Denys Biriukov for their helpful discussions.

## Notes

### Competing Interest Statement

The authors have declared no competing interest.

