## Supplementary Information for "Molecular Dynamics Simulations Unveil the Aggregation Patterns and Salting out of Polyarginines at Zwitterionic POPC Bilayers in Solutions of Various Ionic Strengths"

Page 2.
Table S1. Partial charges of charged groups described with ProsECCo force field used in peptide/bilayer MD simulations.

Page 3.
Figure S1. Non-symmetrized number density profiles for $R_9$ aggregation at POPC bilayers.

**Table S1**: The partial charges in the ProsECCo force field of charged groups in POPC lipid, $R_9$ side chain and termini, $Na^+$, $Ca^{2+}$, and $Cl^-$ ions.

| | POPC lipid | | $R_9$ peptide | | | ions | | |
|---|---|---|---|---|---|---|---|---|
| Scaled groups | $2O(PO_4^-)$ | choline | $Gdm^+$ | $-NH_3^+$ | $-COO^-$ | $Na^+$ | $Ca^{2+}$ | $Cl^-$ |
| Partial charges | -0.655 | +0.75 | +0.75 | +0.75 | -0.75 | +0.75 | +1.50 | -0.75 |

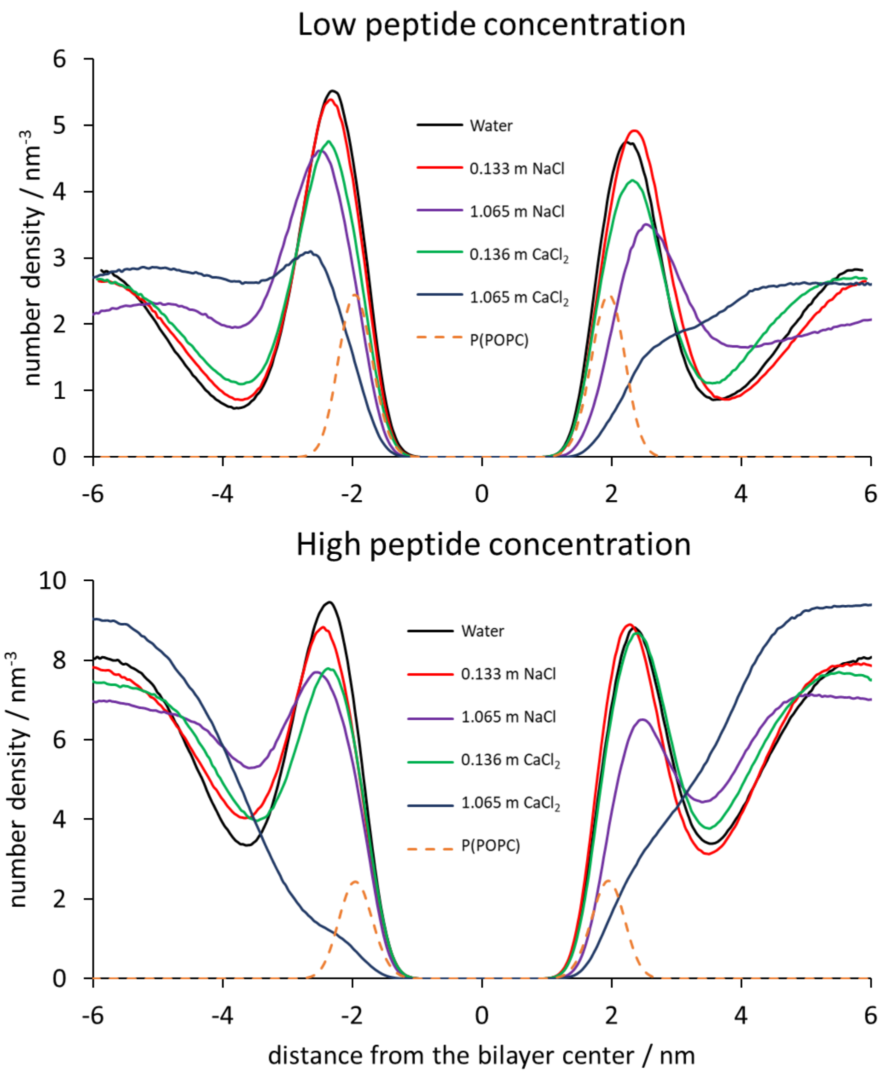


**Fig. S1.** Non-symmetrized number density profiles for peptide center of mass for low $R_9$ concentration (upper panel) and high $R_9$ concentrations (bottom panel) in different systems with respect to the distance from the POPC bilayer center. The number density of POPC phosphorus atoms is shown as the dashed line for the reference system without added salt for low and high peptide concentrations.
